# Vinegar injections can be used safely to control outbreaks of crown-of-thorns starfish (*Acanthaster solaris*) during the peak spawning season

**DOI:** 10.1101/2024.04.02.587823

**Authors:** Pascal Dumas, Amaury Durbano, Bertrand Bourgeois, Hugues Gossuin, Christophe Peignon

**Affiliations:** IRD, UMR 9220 ENTROPIE, BP A5, Nouméa, New Caledonia; IRD, US IMAGO, BP A5, Nouméa, New Caledonia; Aquarium des Lagons de Nouméa, 61 promenade Roger Laroque, 98 800 Nouméa, New Caledonia

**Keywords:** Acanthaster outbreak, vinegar injections, coral reef management

## Abstract

Concerns are mounting over the effects of COTS outbreaks, prompting the need for integrated management strategies. Although direct control methods are short-term and localized, they remain one of the few operational tools that can be easily implemented by local actors. Vinegar injections have recently emerged as highly effective method; however their impact on reproductive behavior remains untested. Here, we investigated the short-term spawning response of mature COTS to double injections of household vinegar. First, COTS abundances and reproductive status were monitored during a massive outbreak affecting New Caledonia’s reefs. In-situ and laboratory experiments were then conducted to determine whether injected COTS would eventually release their eggs and trigger synchronized spawning among mature individuals in close proximity. Our results indicated that injections had no significant effect on spawning behavior, even in densely populated aggregations (>4000 COTS.ha^-1^). In the field, starfish exhibited ripe gonads with high gamete content (up to 35% of body weight) three days after conspecifics were injected. In the laboratory, mature COTS kept with injected, decaying individuals in a confined volume did not expel their gametes after two days. This suggests that vinegar injections could be used at any time, even during peak spawning, without risking synchronized spawning in the affected areas.

## 1. INTRODUCTION

Outbreaks of the coral-eating crown-of-thorns starfish (COTS) present a significant threat to coral reef ecosystems, especially in the Pacific region where concerns are mounting over the increasing frequency and intensity of these once rare and sporadic events (Baird et al. 2013, Pratchett et al. 2017, Milne et al. 2023). New Caledonia, for instance, was once considered relatively unaffected by widespread impacts until the recent decade, when a citizen monitoring program revealed frequent occurrences of spatially localized, yet massive COTS population increases on fringing and mid-shelf reefs (Adjeroud et al. 2018, Dumas et al. 2020). Similar events were documented in Vanuatu, a neighboring archipelago where heightened monitoring efforts raised serious concerns regarding the geographical extent, intensity, and social impact of COTS within coastal communities (Dumas et al. 2014, 2015). In eastern Australia, the Great Barrier Reef has witnessed four extensive outbreaks in the last three decades, with the latest outbreak in 2020 causing significant damage to the northern and central sections of the reef (Pratchett et al. 2021, Bozec et al. 2022).

The recognition of the potential drastic ecological and economic impacts of COTS outbreaks has been gradually increasing within civil society, eventually reaching a point where management efforts are occasionally perceived as inadequate and/or insufficient in certain Pacific island countries (Oremus et al. 2021). As a result, recent studies have underscored the imperative of more comprehensive and integrated strategies, encompassing both theoretical advances necessary to fill knowledge gaps, and more efficient, operational management measures aimed at mitigating the impacts of COTS on the most vulnerable coral ecosystems (Pratchett et al. 2014, 2021, Plagányi et al. 2020, Bonin et al. 2022).

In the Pacific region, coastal communities have long relied on simple, manual collection methods to control COTS populations. Starfish were eventually removed using everyday tools such as knives, spears, hooks, and spearguns, and then killed either in situ or disposed of ashore as part of collaborative (e.g., community-based) efforts (Fraser et al. 2000). However, concerns gradually emerged regarding their efficacy, as evidence suggested that improper practices and failure to consider the COTS reproductive cycle could lead to negligible or even adverse effects, ultimately resulting in increased densities (Lassig 1995, Bos et al. 2013, Messmer et al. 2013). In particular, incorrect timing associated with inadequate handling incurred the risk of massive gamete release, as synchronized spawning can be triggered by chemical communication between ripe individuals in these broadcast spawners (Miller 1989).

The issue also arises with more recent control methods, such as lethal injections of natural acidic compounds (e.g., vinegar, lime juice, powdered citric or oxalic acid), which are progressively replacing and outcompeting chemicals formerly in use (e.g. sodium bisulfate, sodium hypochlorite, ammonium hydroxide, copper sulfate, among others) (e.g. Rivera-Posada & Pratchett 2012, Yamamoto & Otsuka 2013, Moutardier et al. 2015, Buck et al. 2016, Durbano 2017). Double injections of vinegar, in particular, constitute a highly cost-efficient, environmentally friendly option that was increasingly utilized in Vanuatu and New Caledonia during recent outbreaks (Dumas et al. 2021). While their short-term effectiveness is now well established, the physiological and biochemical processes underlying starfish mortality remain incompletely understood. In particular, so far no studies have focused on the potential impacts of lethal injections on COTS spawning behavior during an outbreak.

In this study, we investigated the short-term spawning response in dense aggregations of mature COTS starfish subjected to double injections of vinegar. The research was conducted in the southwestern lagoon of New Caledonia, during a period of massive population outbreaks in 2017 (up to 15,000 COTS per hectare), which severely impacted the highly diverse coral assemblages. This area, covering 5,500 km^2^, has been under surveillance since 2016 as part of a country-wide citizen-monitoring program for COTS (Dumas et al. 2020). First, we surveyed COTS abundances and assessed their reproductive status using the Gonado-somatic index (GSI) during a major outbreak. We then conducted in-situ and laboratory experiments to determine whether injected COTS would eventually release their eggs and trigger synchronized spawning among mature individuals in close proximity.

## 2. MATERIAL & METHODS

### 2.1 COTS census and collection

We carried out quantitative COTS verification on the fringing reefs of the Vua islet, an isolated patch reef located 45 km south-east of Nouméa where emergence of COTS had been reported by a snorkeler (Fig. 1). The area was divided into five consecutive zones based on a preliminary visual assessment of COTS presence. In each zone, COTS were surveyed by a team of two snorkelers swimming parallel to the reef edge during the day. Abundances were estimated using standardized ten-minute swims. The depth range was 1–5 m, with 5–10 m between observers to avoid overlap. A handheld GarminTM Map60Cx GPS device in an underwater housing recorded the position of the timed swims.

**Figure 1.**
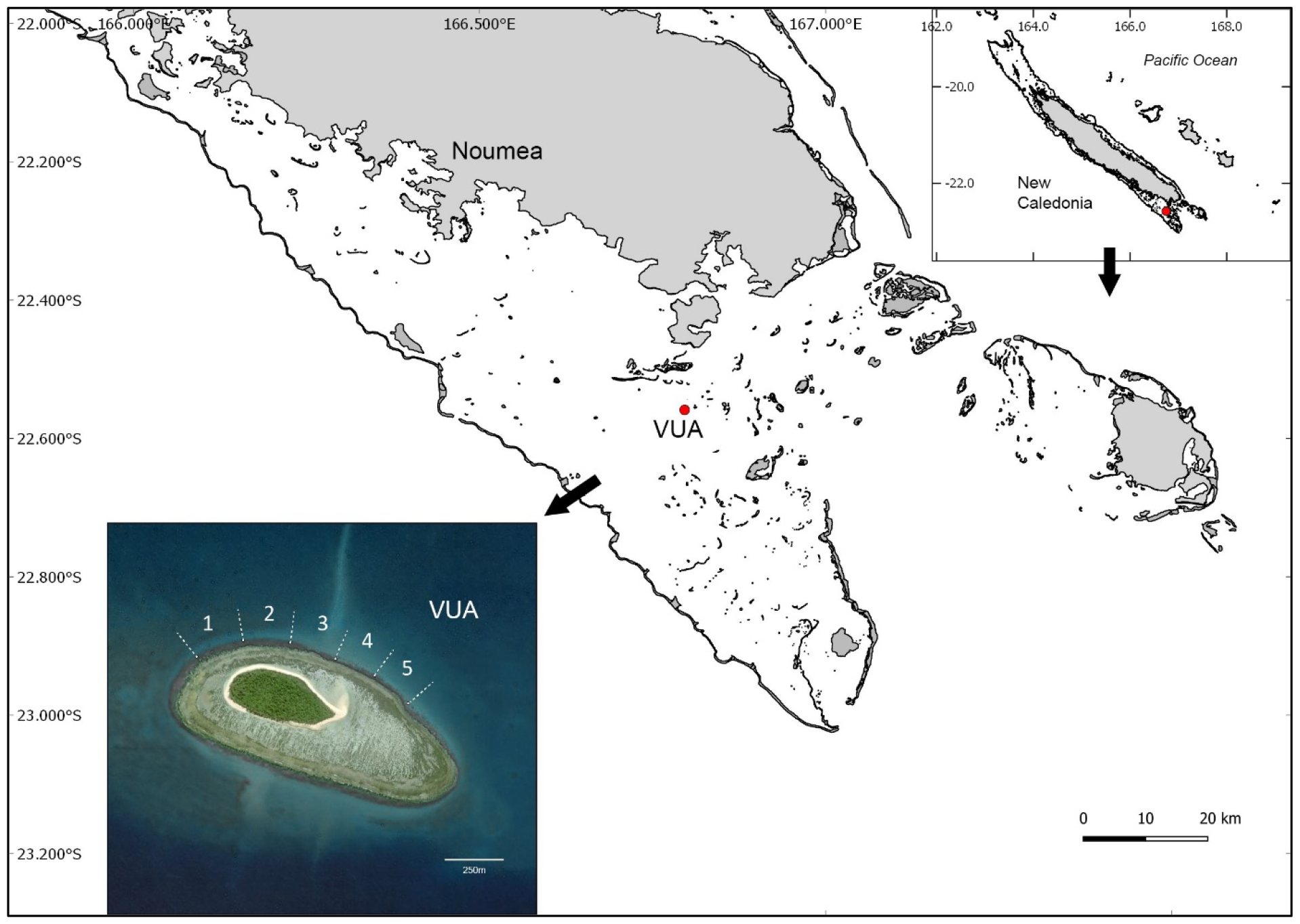
Location of the study area in the southwestern lagoon of New Caledonia.

Fifty specimens were randomly collected from the zone with highest abundances (zone 4) immediately after the census. We targeted large specimens (diameter >30cm) that had reached sexual maturity and appeared to be in good condition. Live COTS were carefully transported to the IRD facilities in containers filled with regularly refreshed seawater, then stored in two flow-through 3000 l raceways with constant filtered seawater inflow for the duration of the experiments.

### 2.2 In situ injection experiment

#### Controls

Fifteen specimens (*controls* hereafter) were dissected on the day of collection (d0). The size (largest diameter of two perpendicular measurements), wet weight and the total number of arms were recorded for each individual. Following Conand (1983), sex was determined based on the texture and color of the gonads (white for males and orange for females). For each specimen, gonads from three randomly chosen consecutive arms were separated and weighed using an electronic scale to calculate the gonado-somatic index (GSI) (Dumas et al. 2016):

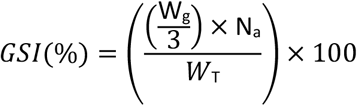

Where:

W_T_: total weight of the specimen (g)

W_g_: cumulated weight for the gonads of three consecutive arms (g);

N_a_: total number of arms

#### In-situ injections

On the next day (d1), lethal injections were carried out in the area where the control samples had been taken (i.e. zone 4). We randomly selected six coral heads with very dense COTS aggregations (≥5 indiv.m^-2^). Each aggregation had its position signaled by a surface buoy attached to the coral substrate. The position of each buoy was recorded by a snorkeler holding the GPS in its underwater housing. In each aggregation, COTS were counted, then one COTS out of two was randomly treated with double injections of (2 × 10 ml) household vinegar in two different areas on the body and left in place, following the method developed by Moutardier et al. (2015). In total, 79 out of 156 COTS were injected.

Three days after the injections (d4), 31 regular (i.e. non-injected) COTS were randomly collected from the same aggregations. We also collected 10 additional individuals at increasing distances from the injection zone: 5 COTS in zone 3 (distance 275m) and 5 COTS in zone 1 (distance 690m). They were immediately transported to the IRD facilities and dissected for GSI calculation, using the same protocol as the control specimens.

### 2.3 Laboratory injection experiment

For this experiment, the collected COTS (n=35) were kept together in the same raceway. All individuals were acclimated for 4 days prior to the start of the experiment, in order to detect any physiological anomalies and to avoid immediate post-capture mortality.

#### Control

At the end of the acclimation phase, five random individuals were removed from the raceway and immediately dissected for GSI calculation using the same protocol (*controls – d0*).

#### Raceway injections

Five random COTS were also treated with double injections of (2 × 10 ml) vinegar in two different areas on the body, using the same protocol as in the in situ experiment. These treated starfish were the left among the other starfish in the raceway. Of the 25 non-injected COTS that remained, ten were dissected on the first day post-exposure to the injected specimens (d1), followed by ten more after two days of exposure (d2). The GSI was calculated for all specimens. The remaining five individuals were kept in the raceway for 30 days with regular visual control of their physiological status.

##### a. Reproductive seasonality

To confirm the timing of the experiments in relation to the peak spawning season in New Caledonia, 68 additional starfish were collected during 17 trips to the area’s reefs between 19/01 and 15/04. The number and location of the COTS actually varied between collection trips, depending on logistical and/or meteorological considerations. The GSI was calculated for each individual using the same protocol; values were then pooled by date.

### 2.4 Data analysis

We used an empirical conversion ratio based upon the average reef surface covered during a typical 10-minute swim to derive estimates of COTS density (expressed in number of COTS per hectare). The area surveyed was calculated by using the linear distance travelled along the reef edge, multiplied by an estimate of the corridor width where observers actually looked for COTS. We used the three-levels abundance/density thresholds recently developed in New Caledonia by Dumas et al. (2020) to classify the COTS populations, where values above five starfish.10 min^−1^/ 100 COTS.ha^-1^ were indicative of an outbreak.

Differences in GSI before and after in situ injections were tested using Student’s T test. ANOVAs were carried out on the GSI to test for the effects of a) distance from the injection zone (in situ experiment); b) time after injections (laboratory experiment). Statistical analyses were performed after logarithmic transformation where necessary to meet the assumption of normality. All analyses were conducted using Statistica 10 package.

## 3. RESULTS

### 3.1 COTS population

Census data confirmed the occurrence of a severe outbreak in the surveyed reef area. Densities ranged from 25 to more than 5000 COTS.ha^-1^, with values well above the threshold considered for outbreak populations in all zones except the last one (zones 1 to 4, average density 2298 COTS.ha^-1^). Densities increased steadily from zone 1 (where the outbreak was first reported) to zone 4 (where the front of the starfish showed maximum densities, e.g. peak value of 5217 COTS.ha^-1^), and then dropped abruptly below the threshold level in zone 5 (average 24.8 COTS.ha^-1^) (Fig. 2).

**Figure 2.**
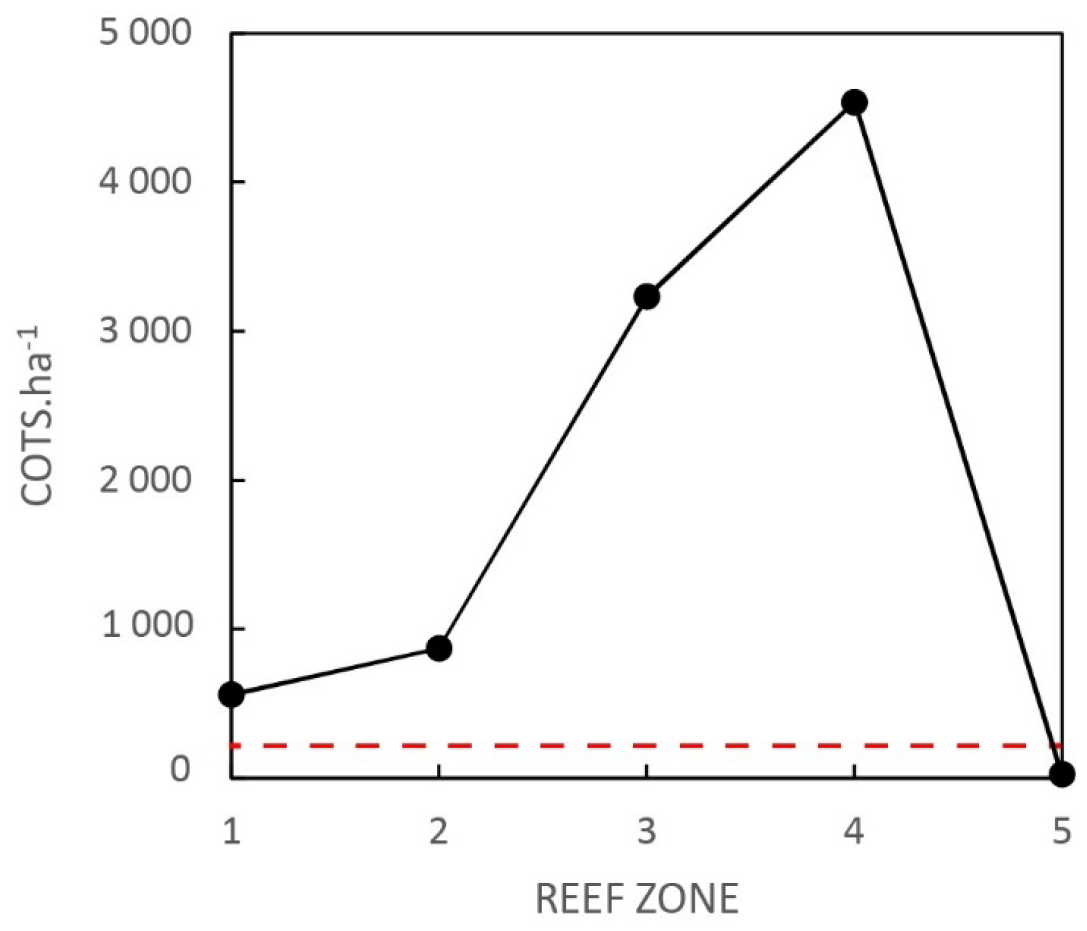
Spatial distribution of COTS abundances in the five zones of the study area before the injection experiment. Means ± SE for number of COTS per 10 minutes swim. Dotted red line: outbreak threshold from Dumas et al. 2020.

In the area, GSI values ranged from 0.2 to 38.9 across the time of the study. The mean GSI increased to reach its maximum at the end of January (20.1), then dropped across February to reach minimum values (<5 overall) from March onwards. Both in situ and laboratory injection experiments were conducted before the drop, i.e. when mature starfish were ready to spawn with GSI close to peak value in the natural populations (average GSI 18.5 ± 1.4 for controls) (Fig. 3).

**Figure 3.**
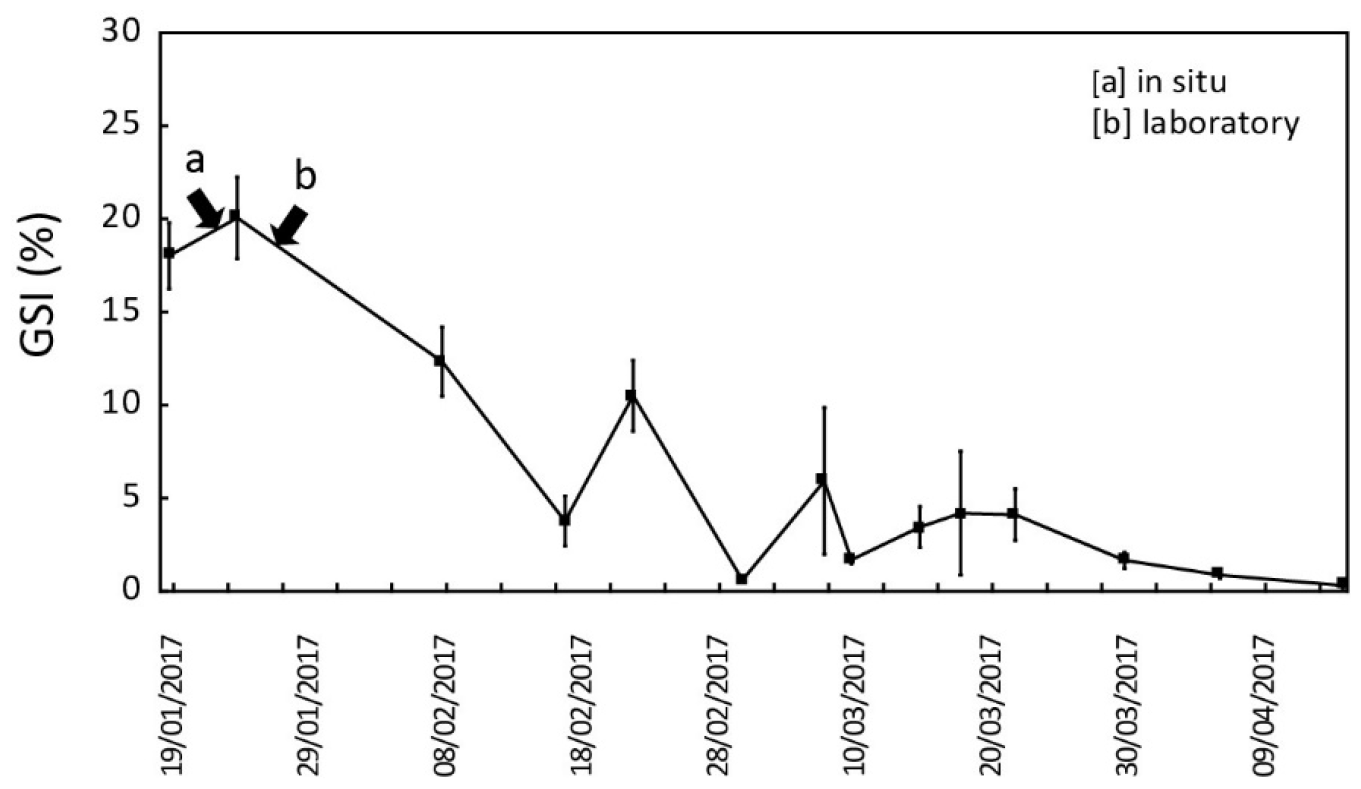
Timing of the injection experiments within the reproductive cycle of *Acanthaster solaris*. Temporal evolution of the gonado-somatic index (GSI) between January and April 2017 in the southwestern lagoon of New Caledonia (means ± SE) and timing of (a) in situ and (b) laboratory injection experiments.

### 3.2 Injection experiments

During the in situ experiment, the non-injected COTS collected in the area three days after the injection campaign showed no external signs of disease or deterioration. Behavioral responses (including color, tonicity, locomotion) during and after the collection process were normal. Dissections confirmed that all specimens were sexually mature and ready to spawn. Sizes (largest diameter) ranged from 35.5 to 51.5 cm, with individual weight between 1288 and 3842g. COTS exhibited high gamete content (GSI 18.9 ± 1.1) with no abnormalities in the shape, color or texture of the gametes.

No significant effect of vinegar injections were detected in the injection zone: GSI values remained consistently high before and after the injection campaign (mean 18.0 vs. 17.9, t=0.009, p=0.99, N.S). Similarly, GSI exhibited no significant variation from the control as distance from the injection zone increased (ANOVA between zones, F=1.093, p=0.36, N.S) (Fig. 4a).

**Figure 4.**
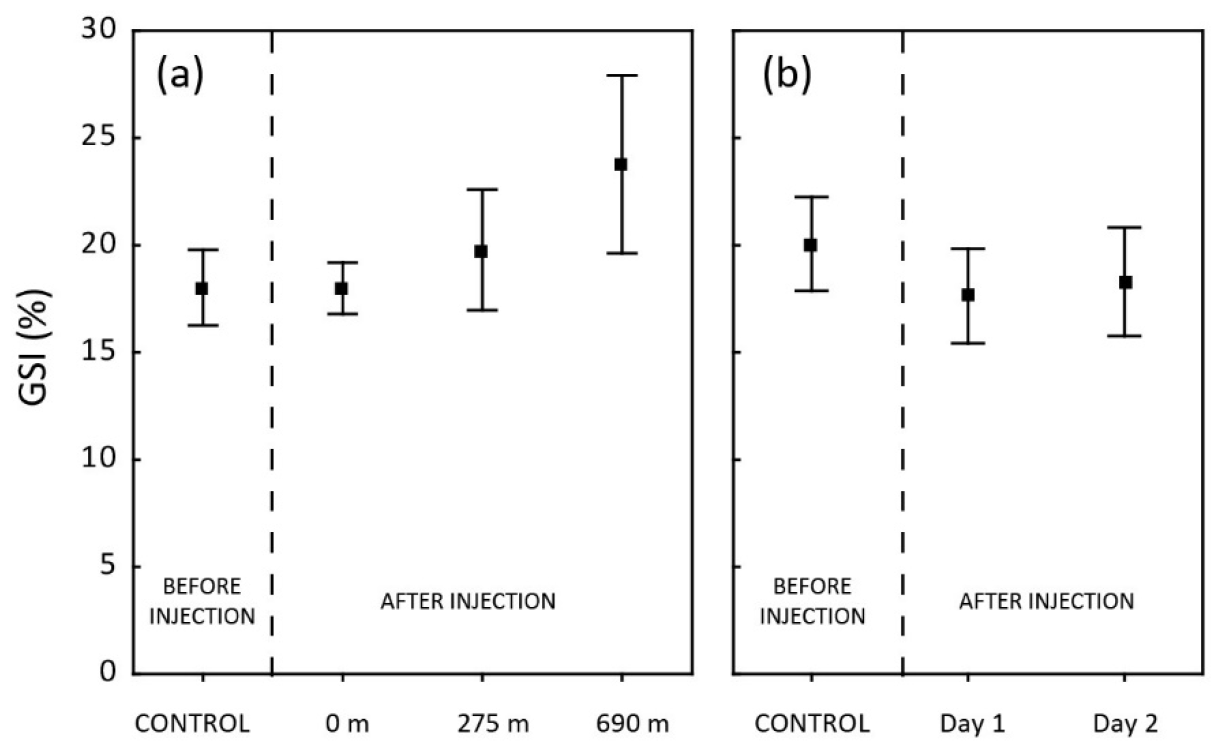
Impact of acidic injections on the spawning of *Acanthaster solaris*. Evolution of GSI before (control) and after injections with respect to (a) distance and (b) time (means ± SE).

Similar results were observed in the laboratory experiment (Fig. 4b). COTS in the raceway ranged from 37 to 50 cm in diameter (928 to 3118 g). All starfish were sexually mature with high gamete content (GSI 18.4 ± 1.4) at the beginning of the experiment, and these values remained consistently stable throughout the 3-day exposure period to the injected conspecifics (ANOVA before/1 day after/2 days after exposure, F=0.197, p=0.82, N.S.) (Table 1).

**Table 1.** Influence of acidic injections on the gonado-somatic index (GSI). Results of ANOVAs on GSI for distance and time factor in a) in situ experiment and b) laboratory experiment (*N*.*S*.: *not significant*).

|  | df | F | p |  |
| --- | --- | --- | --- | --- |
| **a. In situ experiment** |  |  |  |  |
| Distance | 3 | 1.093 | 0.36 | (N.S.) |
| **b. Laboratory experiment** |  |  |  |  |
| Time | 2 | 0.197 | 0.82 | (N.S.) |

## 4. DISCUSSION

Despite extensive scientific research over the past half-century - a simple search for *Acanthaster* or *crown-of-thorns* on the Web of Science currently returns over 1000 scientific articles -, significant gaps persist in our understanding of crown-of-thorns starfish biology and ecology (Pratchett et al. 2014, 2021). Beyond theoretical considerations, this severely hampers our ability to effectively predict and manage sporadic population outbreaks, and prevent widespread coral degradation at the Pacific scale (Mellin et al. 2016, Castro-Sanguino et al. 2023). In this paper, we investigated whether injections of household vinegar, a control method increasingly used in some Pacific island territories, could have detrimental effects by triggering massive spawning among aggregations of mature COTS.

The concept that inadequate control practices may lead to group spawning can be traced back to the early 1970s, when laboratory tests demonstrated chemical communication between COTS during reproduction (Beach et al., 1975). The importance of reproductive triggers/synchronizers was highlighted by experimental studies, along with rare field observations of group spawning eventually induced by the spawning of few individuals (Miller 1989, Babcock & Mundy 1992, Pratchett et al. 2014). Subsequent research suggested that the spawning process may be first initiated by mature males releasing sexual pheromones, along with their gametes, in response to environmental cues (e.g., temperature, photoperiod, salinity, food availability, etc.), which subsequently synchronize females and other males (Babcock et al. 1994, Mercier & Hamel 2009, Caballes & Pratchett 2017). This eventually led to the hypothesis that physical damage inflicted on ripe starfish during manual collection could induce massive spawning, with the leakage of gametes and/or ovarian tissues acting as a spawning cue for conspecifics in close proximity (the “leakage hypothesis”).

The risk is particularly high in the context of community-based cleanups traditionally employed in the Pacific region, where tools used to dislodge COTS from the reef/coral matrix often damage their arms, which contain the gonads. Village clean-ups are generally organized ’on the fly’ when COTS densities reach levels that interfere with daily activities such as reef fishing, invertebrate collection or swimming, without regard to the reproductive cycle of COTS. This was observed during a major village-based cleanup effort conducted in the fringing reefs of Santo (Vanuatu), where locals removed approximately 13,000 starfish during the peak spawning period using knives, spears, hooks, etc. Most starfish exhibited missing arms, tears and perforations, or were cut into pieces during the collection process. Results showed that the gonado-somatic index (GSI) measured at the end of the removals significantly dropped in the cleaned areas compared to the adjacent reefs, providing the first field evidence that traditional removal practices could trigger gamete release (Dumas et al. 2016).

Vinegar injections have gradually emerged as a highly credible control alternative for COTS, combining high effectiveness (e.g. 100% mortality within 12-24 hours for double injections of 10ml, Moutardier et al. 2015) with numerous logistical and cost advantages over more traditional methods (KBRF 2012, Rivera-Posada et al. 2013, Buck et al. 2016, Boström-Einarsson et al. 2018). Initially developed and tested in Vanuatu where it is now routinely used by the National Fisheries Department in support of coastal communities, the double injection method has recently been trialed in New Caledonia as part of a pilot project following the latest wave of outbreaks affecting coral reefs since 2015. One potential benefit stems from the assumption that the needles and injection process do not significantly impair the physical integrity of the starfish, potentially reducing (or even eliminating) the risk of massive gamete ejection associated with manual collection methods. To our knowledge, this study is the first attempt to test this hypothesis in real conditions during an actual outbreak.

Here we used the gonado-somatic index (RGS) as a simple, yet robust proxy for assessing the reproductive status and seasonality of COTS (Conand 1983, Babcock et al. 2016). In New Caledonia, COTS reproduction is relatively condensed in the year compared to lower latitudes, with optimal spawning season concentrated around the end of the austral summer when water temperature reaches 26 °C (Hue et al. 2020). We specifically positioned the study during this critical period, i.e. just before the peak spawning when ripe COTS were most susceptible to synchronized spawning. The temporal patterns of RGS were consistent with the previous findings of Conand (1984), where the highest values (15-20) were reached in January-February, shortly preceding a sudden decline indicative of gamete release in males and females.

Overall, our results emphasize that vinegar injections had no significant impact on the spawning behavior of COTS. Anatomical observations (dissections) and RGS analyses for both in situ and laboratory experiments confirmed that even in densely populated aggregations, mature individuals in close proximity to those subjected to lethal injections did not expel their gametes. In the field, specimens collected from our six random patches with density ≥ 5 COTS.m^-2^ still exhibited non-modified, ripe gonads with high gamete content (up to 35% of total body weight) three days after conspecifics were injected. This was observed equally in male and female starfish. In contrast, the classic signs of decomposition (including discoloration and ulceration of the body wall, tears, loss of arms), as well as the presence of numerous feeding fish, were observed in specimens that were treated by the double vinegar injections. This was further confirmed in even more restrictive laboratory conditions, where ripe COTS were kept together with injected, decaying individuals in a very confined volume (∼3 m^3^) with limited water renewal. Analyses revealed that all non-injected individuals retained mature and apparently functional gonads with high gamete content (max 37.8% of total body weight) up to 2 days after conspecific injection, with no evidence of spawning.

These results are particularly promising as they remove one of the major uncertainties associated with manual methods. In the absence of direct manipulation of the COTS, the injection process does not cause major physical damage, and the perforations are likely not sufficient to cause significant leakage of gametes and associated pheromones. While the timing of the control campaigns remains essential to optimize outcomes - ideally by anticipating the reproduction season and thus the production of new cohorts - our findings show that it would be possible to carry out the injection campaigns at any time, including at the peak of the breeding season, without risking massive, synchronized spawning in the affected areas. This is all the more relevant as the exact time of reproduction can vary considerably according to latitude and specific locations, which makes it difficult to frame control campaigns within the COTS reproductive cycle in areas where this information is lacking (Yasuda et al. 2010). Of course, it is important to bear in mind the limitations of this study, in particular the relatively short post-injection period (3 days in situ, 2 days under experimental conditions) during which the gonads of the specimens were monitored. However, the hypothesis of an even later response (>3 days) to the injections seems very unlikely, as the available data indicate that spawning typically occurs within minutes to 1-2 hours after the initiation of the process (Babcock & Mundy 1992, Caballes & Pratchett 2017). This was also observed in the laboratory trial, where the last five COTS did not show subsequent reaction during the 30 days after injection, when the animals were kept for observation.

Field observations during various pilot campaigns in New Caledonia and Vanuatu have confirmed that double injections of vinegar are highly effective in culling COTS, particularly when extreme densities (e.g. > 5000 COTS.ha^-1^, Buttin 2018, Dumas et al. 2022) place considerable human, financial and logistical constraints on more conventional manual collection methods. However, there are still uncertainties about the physiological processes involved in the death of the COTS starfish exposed to vinegar. It is hypothesised that the generalized decay observed after injections may be caused by the inability of COTS to regulate their pH, with a subsequent immunological response to the damaged tissues leading to the death of the injected starfish. Elevated pH may also promote the selective growth of pathogens favored by acidic environments, as observed with TCBS injections which induce rapid death by generalized bacterial infection and/or allergic reaction (Rivera-Posada et al. 2011, 2012, Grand et al. 2014, Rivera-Posada & Owens 2014). Preliminary experiments on gametes after vinegar injection have also suggested the presence of several markers associated with cellular degeneration (Durbano 2017), prompting questions about their viability should they be released after injection by decaying starfish. Further research is then needed to validate these hypotheses and to investigate the cellular and subcellular processes leading to death specifically in vinegar.

Longer-term, integrated management strategies are needed to more effectively address the COTS phenomenon and associated decline of coral cover on a large (Indo-Pacific) scale. Although control methods are inherently short-term and localized solutions, they remain one of the few operational levers available that can be easily implemented by local actors including coastal communities and managers. Given their high cost-effectiveness, minimal environmental and eclogical impact, and logistical ease, vinegar injections emerge as one of the most promising control methods for countries affected by COTS.

## Acknowledgements

The authors express their warmest thanks to the technical staff of the IRD Noumea for technical assistance: Miguel Clarque, Samule Tereua and Philippe Naudin. We are also very grateful to the staff of the Aquarium des Lagons for providing logistical support, in particular to Hugues Gossuin, Sylvain Govan and Olivier Chateau. This work was conducted within the framework of IRD COTS research programme in New Caledonia, including the OREANET initiative (www.oreanet.ird.nc) funded by the Government of New Caledonia and the French Fonds Pacifique.

